# Nontypeable *Haemophilus influenzae* infection impedes *Pseudomonas aeruginosa* colonization and persistence in mouse respiratory tract

**DOI:** 10.1101/2021.08.05.455360

**Authors:** Natalie Lindgren, Lea Novak, Benjamin C. Hunt, Melissa S. McDaniel, W. Edward Swords

## Abstract

Patients with cystic fibrosis (CF) experience lifelong respiratory infections which are a significant cause of morbidity and mortality. These infections are polymicrobial in nature, and the predominant bacterial species undergo a predictable series of changes as patients age. Young patients have populations dominated by opportunists that are typically found within the microbiome of the human nasopharynx, such as nontypeable *Haemophilus influenzae* (NTHi); these are eventually supplanted and the population within the CF lung is later dominated by pathogens such as *Pseudomonas aeruginosa* (*Pa*). In this study, we investigated how initial colonization with NTHi impacts colonization and persistence of *Pa* in the respiratory tract.

Analysis of polymicrobial biofilms *in vitro* by confocal microscopy revealed that NTHi promoted greater levels of *Pa* biofilm volume and diffusion. However, sequential respiratory infection of mice with NTHi followed by *Pa* resulted in significantly lower *Pa* as compared to infection with *Pa* alone. Coinfected mice also had reduced airway tissue damage and lower levels of inflammatory cytokines as compared with *Pa* infected mice. Similar results were observed after instillation of heat-inactivated NTHi bacteria or purified NTHi lipooligosaccharide (LOS) endotoxin prior to *Pa* introduction. Based on these results, we conclude that NTHi significantly reduces susceptibility to subsequent *Pa* infection, most likely due to priming of host innate immunity rather than a direct competitive interaction between species. These findings have potential significance with regard to therapeutic management of early life infections in patients with CF.

## INTRODUCTION

Cystic fibrosis (CF) is an autosomal recessive genetic illness that causes dysfunctional ionic transport at epithelial surfaces, with concomitant impacts on mucociliary defenses and clearance in the lungs. As a consequence, patients with CF experience lifelong infections of the airway mucosa that cause chronic inflammation and epithelial damage, among other sequelae (1–3). Despite significant therapeutic gains that have significantly increased average lifespan for CF patients, airway infections and associated respiratory complications remain a leading cause of morbidity and mortality in patients with CF disease (4). The microbial populations within the CF lung are complex, polymicrobial communities that undergo dynamic changes that correlate with patients’ age and overall health, and can be a determinant of disease severity (3, 5–7). Despite years of study of opportunistic infections in the context of CF, there remains much to be learned about how specific pathogens and opportunists impact the course and severity of disease (6–9). Nontypeable *Haemophilus influenzae* (NTHi) is a non-encapsulated, Gram-negative opportunist that normally resides in the human nasopharynx and upper airways as part of the normal flora. (10–12). Although typically benign in healthy individuals, NTHi is among the most common bacterial species isolated from CF patients during the first year of life and is thus thought of as a common early-stage pathogen (13–15). As with other mucosal airway infections, NTHi persists in the lung within biofilm communities (16, 17). Like other mucosal pathogens, NTHi produces lipooligosaccharide (LOS) endotoxins with truncated carbohydrate regions (18). Like most endotoxins, the NTHi LOS elicits host inflammation via Toll-like receptor 4 and is likely an integral component for symptomatic response to NTHi infection (19, 20).

As CF patients age, the bacterial population diversity typically declines, and the microbial population in the lung becomes dominated with late-stage pathogens, such as *Pseudomonas aeruginosa* (*Pa*) (8, 9, 21, 22). *Pa* is a Gram-negative opportunistic pathogen that has a high level of inherent resistance to antimicrobials and host immune effectors and is therefore difficult to treat (23). To adapt to the CF airway environment, *Pa* populations typically undergo mucoid conversion to overexpress the exopolysaccharide alginate, which is a component of *Pa* biofilms (24, 25). Mucoid conversion of *Pa* is associated with increased antibiotic resistance, persistent inflammation, and overall worse clinical outcomes (26). Initial *Pa* infections, which may take place within the first year of life, frequently become chronic after mucoid conversion (19, 20). While the role of *Pa* in end-stage CF disease has been well established, less has been done to establish the role of *Pa* in early-stage CF disease, particularly the ages in which NTHi colonization peaks in frequency (4, 9).

Bacteria reside in the respiratory system as a complex, evolving polymicrobial community in which individual organisms may interact with one another in a synergistic, antagonistic, or null fashion (11, 27–29). Changes in these communities as different species, or strains within species, interact are thought to greatly influence antibiotic susceptibility, immune evasion, and mutations for chronic colonization, all influencing patient outcomes (8, 30). To fully understand how specific species contribute to disease pathogenesis, we must characterize community members in concert, modeling their interactions after the environment in which they naturally reside, rather than as individual species (5, 8, 31). In this study, we aimed to assess the polymicrobial interaction between NTHi and *Pa* both *in vitro* and *in vivo*, using clinically isolated bacterial strains from patients with CF. Mice were intratracheally infected with NTHi and/or mucoid *Pa* (mPA) to produce localized, acute respiratory infections. To model the temporal acquisition of NTHi and *Pa* in CF patients, we tested whether pre-colonization with NTHi influences colonization of *Pa* and began to characterize how this interaction may influence the innate immune response.

## RESULTS

### Nontypeable *H. influenzae* and *P. aeruginosa* form polymicrobial biofilms *in vitro* that support *Pa* growth

In any environment that contains more than one bacterial species, including the CF lungs, organisms may interact with one another synergistically, competitively, or null (4)(5). To begin to characterize the polymicrobial interaction between NTHi and *Pa*, we first wanted to evaluate how these two organisms form a polymicrobial biofilm *in vitro*. Since NTHi and *Pa* are typically acquired sequentially over a patient’s lifespan, dual-species static biofilms were sequentially seeded with NTHi followed by *Pa* 12 hours later. These polymicrobial biofilms were grown for a total of 18 hours, while single-species biofilms of NTHi and *Pa* were grown for 12 hours and 6 hours, respectively, at 37°C + 5% CO_2_. Timepoints for growth were selected based on the peak viability of NTHi and *Pa*, as determined by counts of viability over time (Fig. S1A in supplemental material). Utilizing various strains of NTHi, we assessed viable colony counts of *Pa* from sequential dual biofilms compared to single-species *Pa* biofilms. *In vitro*, pre-colonization with NTHi resulted in significant increases in *Pa* growth when both organisms are inoculated at roughly the same amount, suggesting a synergistic relationship (**P<0.01) (Fig. S1B-D).

We then visualized NTHi and *Pa* single- and dual-species fluorescent biofilms with confocal laser scanning microscopy (CLSM) after 18 hours of growth using strains NTHi 86028NP-gfp+ and mPA 08-31-mCherry+. In single-species biofilms, both organisms formed lawn structures with some towers (Fig. 1A and B). Sequentially seeded dual-species biofilms showed a complex lawn with more diffuse *Pa* colony formation that appeared to support *Pa* growth. As expected, the spatial orientation of dual biofilms were stratified, reflecting the sequential introduction of NTHi followed by *Pa*, with a significant lawn structure of NTHi on the bottom (Fig. 1C). Quantification of *Pa* in both single- and dual-species biofilms by BiofilmQ software revealed that the total volume of *Pa* biofilm is greater in dual-species biofilms than in single-species *Pa* biofilms (Fig. 2A). Segmentation within BiofilmQ software allowed for fluorescent images to be divided by 1 μm cubes with each cube being individually tracked. Following segmentation, the mean cube surface is calculated as the number of points between occupied and unoccupied volume in cube. The calculated mean cube surface showed that dual biofilms have more *Pa* than single-species *Pa* biofilms, recapitulating the results seen with total biofilm volume and verifying the accuracy of cubing (Fig. 2B). This segmentation technique was used for further 3D visualization of single- and dual-species biofilms of *Pa.* Segmentation shows that there is more diffuse local density of *Pa* associated with dual-species (Fig. 2C) rather than single-species (Fig. 2D) growth. Overall, quantification with BiofilmQ and viable colony counts from co-culture suggests that *in vitro*, NTHi positively influences *Pa* growth, producing greater biofilm volume and more dispersed spatial distribution when grown together compared to a single-species *Pa* biofilm.

**Figure 1:**
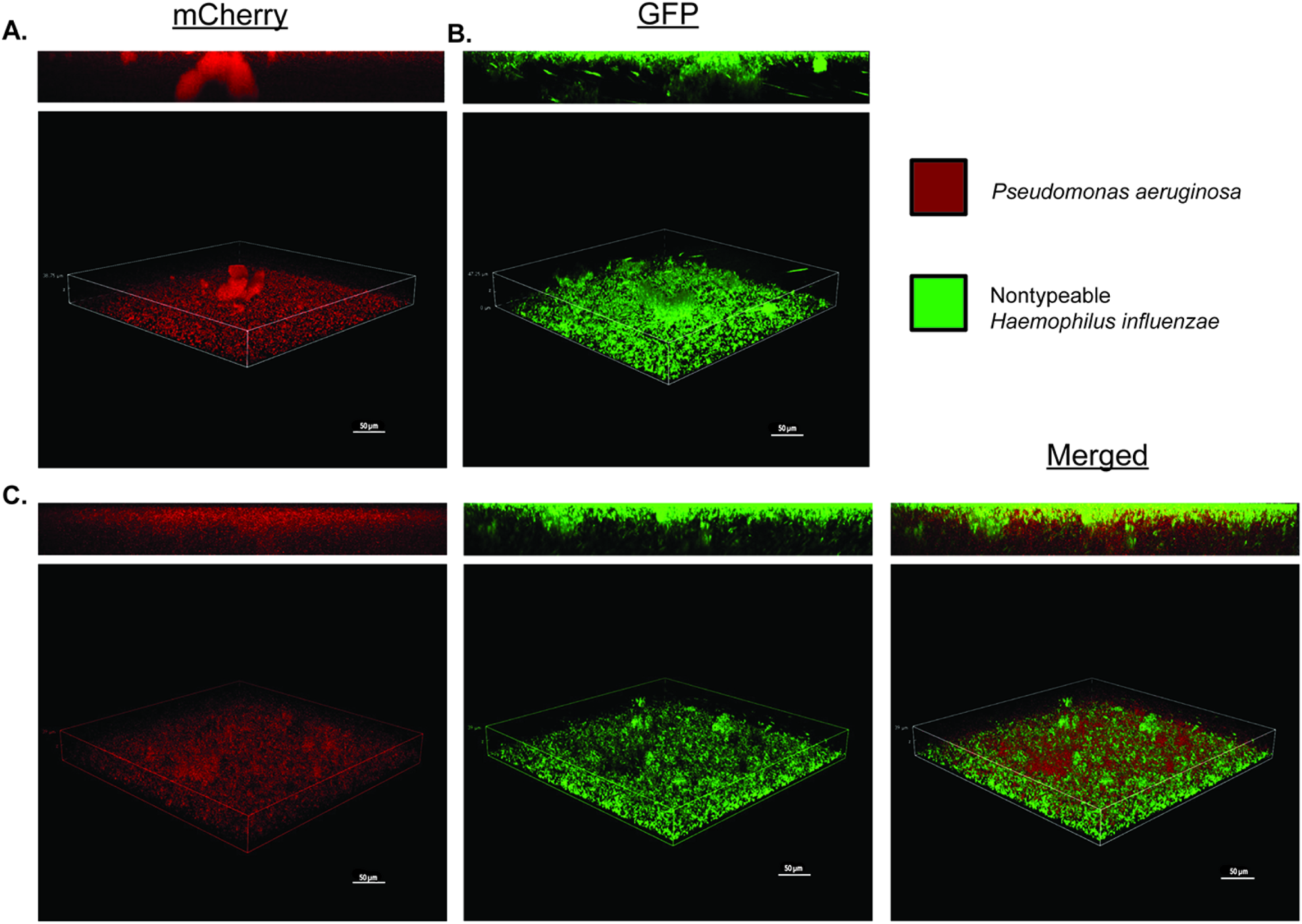
**Nontypeable *H. influenzae* and *P. aeruginosa* form polymicrobial biofilms *in vitro*.** Biofilms on abiotic glass surface of A) *P. aeruginosa* mPA 08-31 (pUCP19+mCherry+) B) Nontypeable *H. influenzae* 86-028NP (PGM1.1+gfp) and C) NTHi and *Pa*. Inoculation of NTHi preceded *Pa* by 12 hours and both organisms grew for additional 6 hours at 37 °C + 5% CO_2_ (18 hours total). Imaged at 40X magnification. Scale bar represents 50 μm.

**Figure 2:**
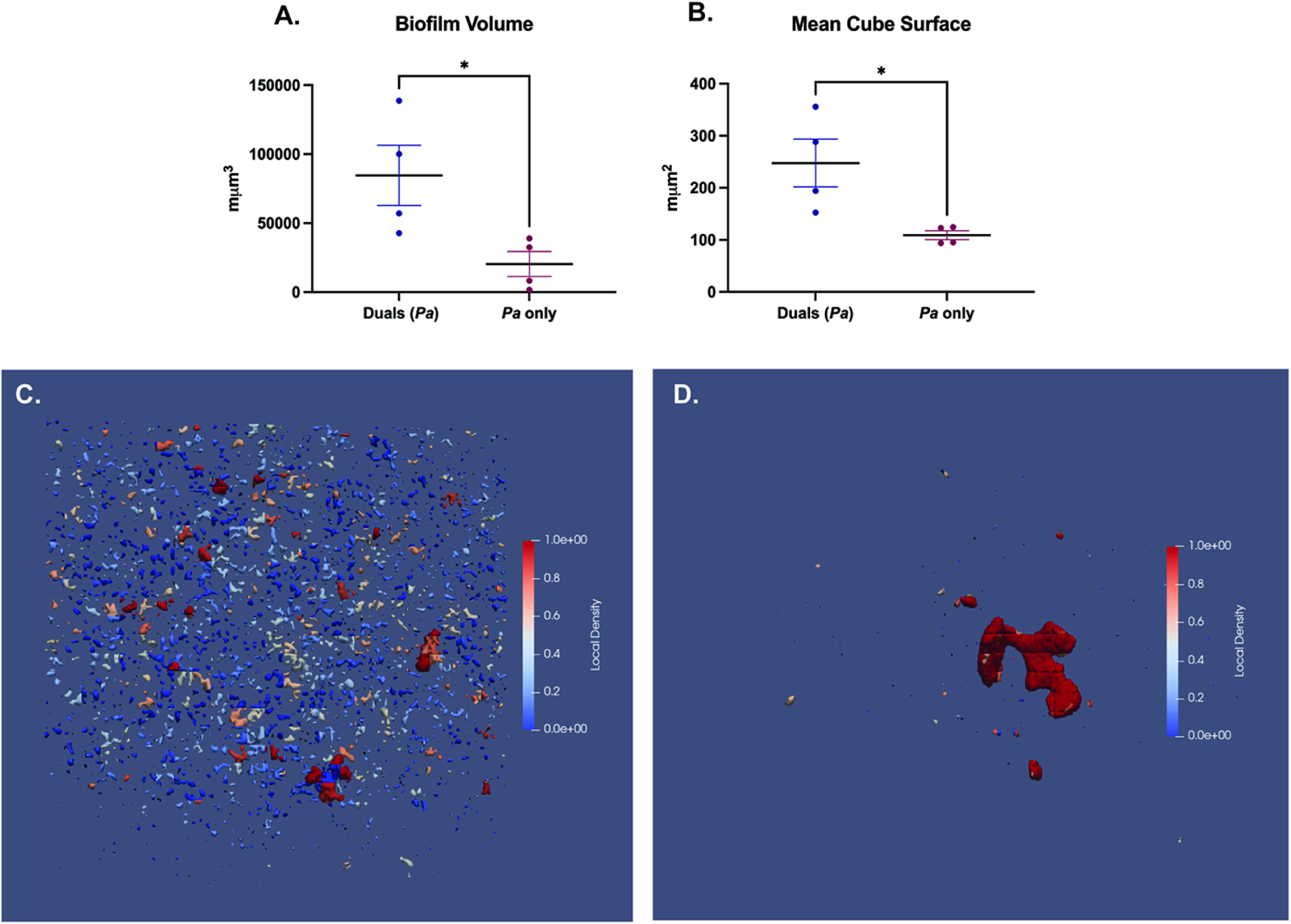
**Quantification of nontypeable *H. influenzae* and *P. aeruginosa* polymicrobial biofilms.** Bacterial density within fluorescent biofilms was quantified using BiofilmQ software to measure A) biofilm volume of *Pa* in dual and single-species biofilms and B) mean cube surface of *Pa* quantified as 1 cube = 1 μm. Cubing can be visualized as local density of *Pa* in both C) polymicrobial NTHi/*Pa* biofilm and D) *Pa* biofilm. Mean ± SEM. (N = 4). Mann-Whitney test, two-tailed, *P<0.05.

### Nontypeable *H. influenzae* infection decreases bacterial burden of *P. aeruginosa* in mouse lung

While our *in vitro* results suggest that NTHi and *Pa* may positively interact in polymicrobial biofilms, we aimed to evaluate how each species is impacted during polymicrobial infection within the host. Through serial passage, we induced antibiotic resistance in a CF sputum-derived clinical isolate of NTHi, HI-1, developing a model strain for acute infection (NTHi HI-1r) that can be differentially plated in the presence of *Pa*. To model the sequential acquisition of an early life pathogen (NTHi) followed by a later-stage pathogen (*Pa*) that typically ensues as CF patients age, BALB/cJ mice were intratracheally infected with NTHi HI-1r (inoculum ~10^8^ CFU/mouse) followed by a mucoid *P. aeruginosa* isolate, mPA 08-31 (inoculum ~10^7^ CFU/mouse) 24 hours later (Fig. S2A). Both organisms colonized the lungs and were detectable by viable plate counting of lung homogenate 48 hours post-infection. However, colonization of

*Pa* was significantly impeded after sequential introduction with NTHi compared to a *Pa* single-infection (**P<0.01) (Fig. 3A). As this contrast the *in vitro* findings, this result supports the importance of studying polymicrobial interactions within a host to accurately model human diseases. To determine the importance of temporal spacing in this interaction, we also infected mice with both organisms concurrently (inoculum ~10^7^ CFU/mL of each) (Fig. S2B). When we compared the bacterial burden of *Pa* in the lungs after a single-infection, a sequential dual-infection, and a concurrent dual-infection, we saw that mice sequentially infected with NTHi followed by *Pa* showed significant reduction in *Pa* abundance compared to a *Pa* only infection (*P<0.05). Additionally, sequential dual-infection resulted in less *Pa* in lung homogenate than concurrent dual-infection (****P<0.0001) (Fig. 3B). This suggests that the competitive interaction between NTHi and *Pa* is dependent on time of introduction, and *Pa* colonization is only impeded following sequential infection. Histopathological analysis of hematoxylin and eosin (H&E) stained lung sections indicated immune cell infiltration and inflammation 48 hours after infection of both NTHi and *Pa* infection (Fig. 3C). H&E stained slides were graded for severity of lung injury as indicated by neutrophil infiltration of alveoli, peri-vascular inflammation and peri-bronchial inflammation. At 48-hours post-infection, where a high burden of both NTHi and *Pa* were present, there was moderate tissue damage for all infection groups with no significant difference between groups. All infections resulted in significantly more severe tissue damage than a PBS vehicle control (****P<0.0001) (Fig. 3D). Immune cell infiltration was expressed by total cell infiltrates counted in the bronchoalveolar lavage fluid (BALF) after cytospin preparation and differential staining. As expected from histological results, all infection groups showed significantly more total cell counts, as well as PMN infiltrates in those counts, than uninfected controls with no significant difference between infection groups (*P<0.05, **P<0.01, ***P<0.001, ****P<0.0001) (Fig. 3E-F). Weight loss per animal was evaluated over the course of infection and weight loss increased compared to uninfected controls (Fig. S3A). Taken together, these results suggests that *in vivo*, NTHi impedes the bacterial burden of *Pa* in the airways when introduced sequentially, but not concurrently, and infection with either organism results in respiratory tissue damage with significant inflammatory response.

**Figure 3:**
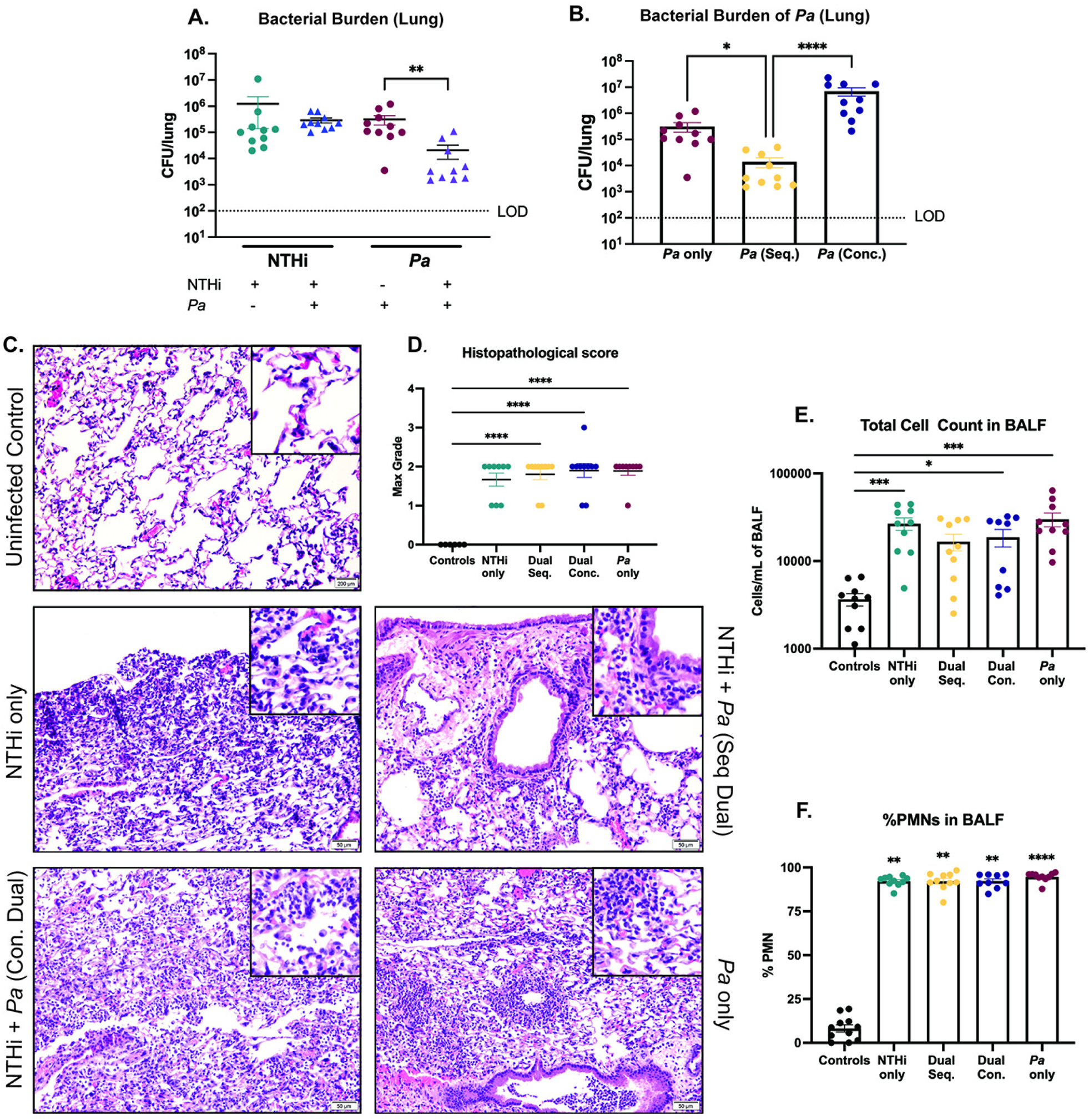
**Nontypeable *H. influenzae* infection significantly decreases *P. aeruginosa* colonization/persistence in mouse lung.** BALBc/J mice were intratracheally infected with nontypeable *H. influenzae* (NTHi Hi-1r) either 24 hours prior to introduction of mucoid *P. aeruginosa* (mPA 08-31) in a sequential fashion, or were introduced at the same time (concurrently) and assessed 48 h post-infection. A) Bacterial counts of NTHi and *Pa* in the lung homogenate enumerated by plate count. B) Bacterial counts of *Pa* in the lung homogenate comparing sequential and concurrent infection. Severity of disease was indicated by C) H&E stained lung sections (200 x magnification) and D) semi-quantitative grading by a pathologist (L.N.). E) Total cell counts in BALF quantified by differential cell counts. F) Percentage of PMNs in the BALF. Mean ± SEM. (N = 10). LOD = limit of detection. Kruskal-Wallis with Dunn’s multiple comparison test, * P<0.05, ** P<0.01, *** P<0.001, **** P<0.0001.

### Impacts of nontypeable *H. influenzae* on *P. aeruginosa* are not strain-specific

Our coinfection results indicate that a clinical isolate of NTHi, HI-1r, significantly inhibits the bacterial burden of mucoid *P. aeruginosa* isolate mPA 08-31 in the airways. To determine if the impact of NTHi on *Pa in vivo* is strain specific, we induced antibiotic resistance in laboratory strain NTHi 86-028NP to develop strain NTHi 86r, allowing for differential plating of NTHi and *Pa*. Following the sequential infection schematic previously described, we intratracheally introduced NTHi 86-028NPr (inoculum ~10^8^ CFU/mouse) into the airways of BALBc/J mice followed by introduction of mPA 08-31 (inoculum ~10^7^ CFU/mouse) 24 hours later (Fig. S2C). Unlike the positive interaction between NTHi and *Pa* seen in our polymicrobial biofilms with the parent strain of NTHi 86r (NTHi 86-028NP), in the airways, viable colony counts of *Pa* were significantly decreased after pre-introduction with NTHi 86r compared to a *Pa* single-infection (*P<0.05) (Fig. 4A). This finding validates that the contrasting, synergistic interaction noted between NTHi 86-028NP and mPA 08-31 seen *in vitro* (Figure 1–2) are dependent on the lack of host response rather than strain specificity of NTHi. Mucoid conversion of *Pa* is a common consequence of long-term colonization (22, 27, 32). For young patients, the initial colonizing *Pa* strain is nonmucoid for a variable length of time, not yet producing the alginate component that is heavily associated with persistent infection (22). To extend the impact of this phenotype to the minority of patients colonized with *Pa* before mucoid conversion, we tested whether pre-introduction with NTHi impedes the abundance of non-mucoid *Pa.* To do this, we performed sequential infections with the clinical NTHi HI-1r followed by a non-mucoid *Pa* (PA FRD1*mucA+).* Viable colony counts from lung homogenate revealed that non-mucoid isogenic mutants of *Pa,* in addition to the mucoid strain previously evaluated, are significantly diminished after pre-introduction with NTHi compared to *Pa* alone (*P<0.05) (Fig. 4B). Weight loss per animal was evaluated over the course of infection and weight loss increased compared to uninfected controls (Fig. S3B-C). These results show that NTHi infection significantly reduces the colonization/persistence of *Pa*, regardless of the specific strain of NTHi or the mucoidy status of *Pa*.

**Figure 4:**
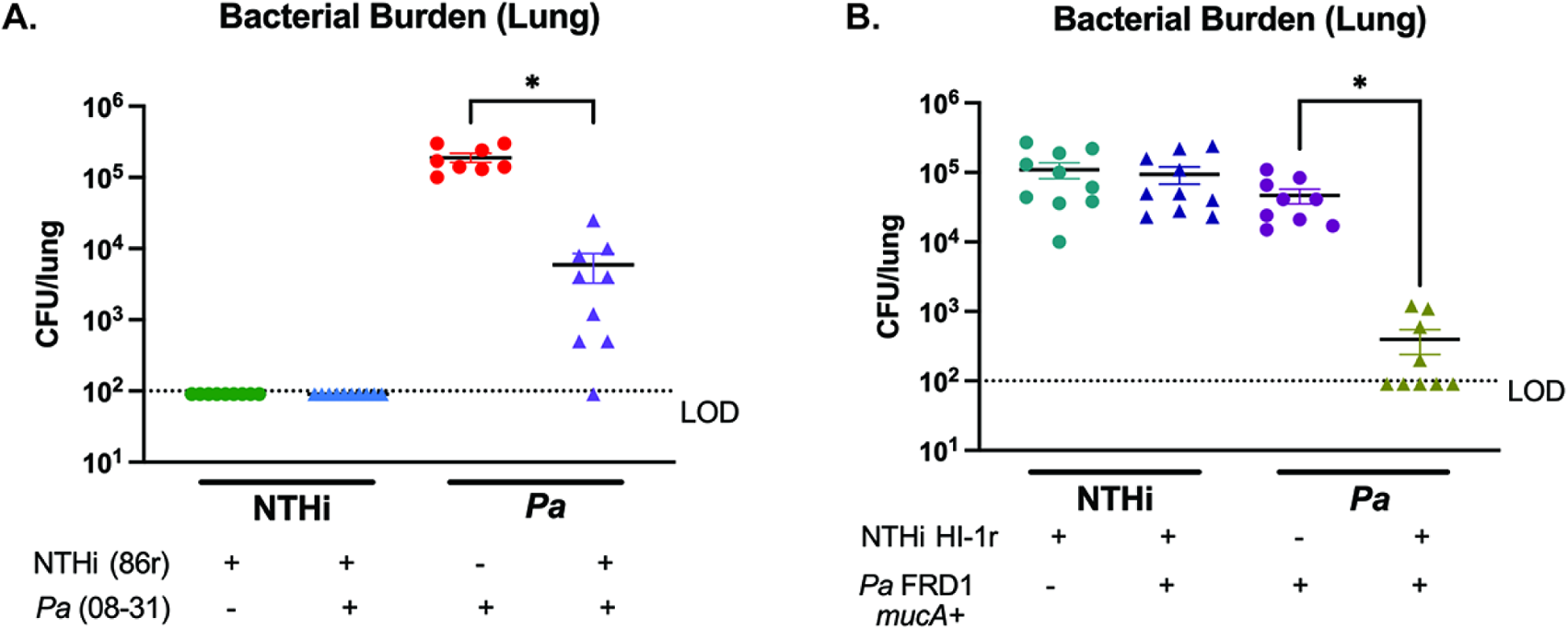
**Nontypeable *H. influenzae* impact on subsequent *P. aeruginosa* infection is strain independent.** BALBc/J mice were intratracheally infected with spectinomycin-resistant nontypeable *H. influenzae* 86-028NP (NTHi 86r) followed by mucoid *P. aeruginosa* (mPA 08-31). A) Bacterial counts in the lung homogenate were enumerated by viable colony counting 72 hours post-infection. Similarly, intratracheal infections were also performed with spectinomycin-resistant nontypeable *H. influenzae* HI-1 (NTHi HI-1r) and nonmucoid *P. aeruginosa* (PA FRD1*mucA+)*. B) Bacterial counts in the lung homogenate were enumerated by viable colony counting 72 hours post-infection. Mean ± SEM. (N = 10). LOD = limit of detection. Kruskal-Wallis with Dunn’s multiple comparison test, * P<0.05, ** P<0.01, *** P<0.001, **** P<0.0001.

### Infection with nontypeable *H. influenzae* followed by *P. aeruginosa* results in less severe airway tissue damage and less inflammatory cytokine infiltration than *P. aeruginosa* alone

While clinical isolate NTHi Hi-1r persists longer than previously established NTHi strains, such as NTHi 86-028NP (data not shown), this strain still begins to be cleared from the airways of mice around 72 hours post-infection, with viable colony counts below the limit of detection for some animals. We wanted to test if the duration of *Pa* inhibiton in the lungs extended past the optimal timepoint for NTHi viability, when less viable bacteria would be present for inter-species interaction. BALBc/J mice were intratracheally infected with NTHi HI-1r, followed by mPA 08-31 24 hours later and viable colony counts from lung homogenate were assessed after 72 hours post-dual infection, extending the previously established 48 hour timepoint (Fig. S2C). As expected, sequential introduction of NTHi prior to *Pa* significantly reduced colonization of *Pa* compared to a *Pa* alone (***P<0.001) (Fig. 5A). Reflecting the inhibition of *Pa* abundance in viable colony counts during dual-infection, histological grading of tissue damage revealed a significantly decreased damage score in dual-infected animals compared to a *Pa* single-infection (**P<0.01) (Fig. 5B). H&E staining of lung sections of each infection group revealed more severe immune cell infiltration, pleuritis, perivascular inflammation, and peri-bronchial inflammation 72 hours post-infection than was seen at the 48 hour post-infection timepoint (Fig. 5C). Finally, total cell counts in BALF and the percentage of PMN infiltrates in those counts was not different among single or dual-infection groups, but was significantly increased compared to uninfected controls (*P<0.05, **P<0.01, ****P<0.0001) (Fig. 5D-E). Weight loss per animal was evaluated over the course of infection and weight loss increased compared to uninfected controls (Fig. S3D).

**Figure 5:**
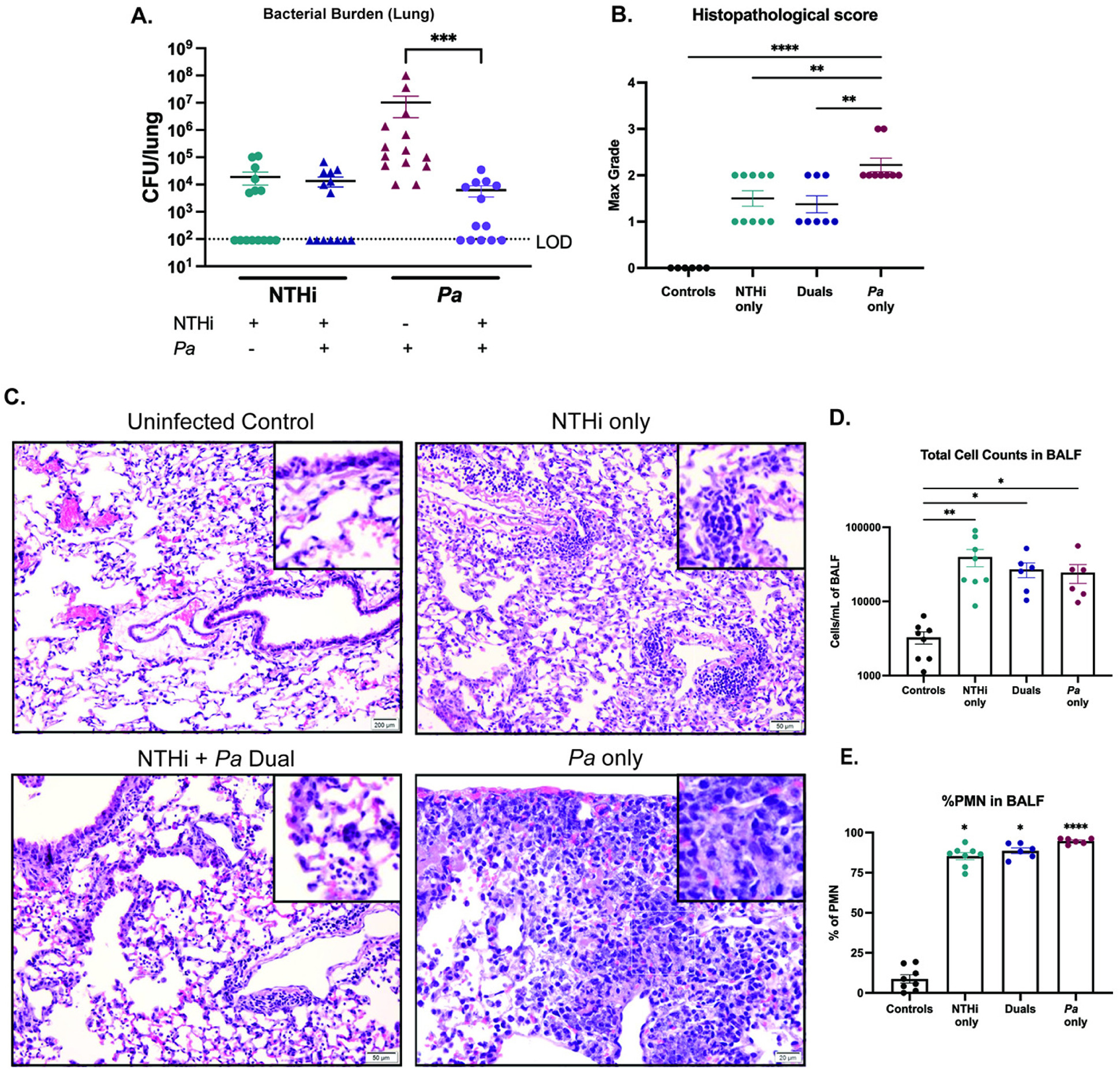
**Nontypeable *H. influenzae* and *P. aeruginosa* dual-infection reduces the severity of airway tissue damage.** BALBc/J mice were intratracheally infected with nontypeable *H. influenzae* (NTHi Hi-1r) 24 hours prior to introduction of mucoid *P. aeruginosa* (mPA 08-31) and assessed 72 hours post-infection. A) Bacterial counts of NTHi and *Pa* in the lung homogenate enumerated by viable colony counting. (N = 13-15). Severity of disease was indicated by B) semi-quantitative grading by a pathologist (L.N.) and C) H&E stained lung sections (200 x magnification) (N = 6-10). D) Total cell counts in BALF quantified by differential counting. (N = 6-8). E) Percentage of PMNs in the BALF (N = 6-9). Mean ± SEM. LOD = limit of detection. Kruskal-Wallis with Dunn’s multiple comparison test, * P<0.05, ** P<0.01, *** P<0.001, **** P<0.0001.

To evaluate the host immune response following NTHi and *Pa* single or dual-infections, we measured cytokine and chemokine levels in BALF with DuoSet ELISA kits designed to detect both pro-inflammatory and anti-inflammatory markers including MCP-1, TNF-α, IL-10, IL-1α, IL-1β, or IL-6. Several of these well-established pro-inflammatory markers, including MCP-1, TNF-α, and the anti-inflammatory marker IL-10, were significantly higher after *Pa* single-infection compared to a dual-infection with both NTHi and *Pa* (****P<0.0001), (Fig. 6A-C). While IL-1α, IL-1β and IL-6 levels were not different between single and dual-infected groups at 72 hours post-infection, dual-infection resulted in less increase of IL-1α and IL-β over uninfected controls than single-species infection (Fig. 6D-F). Taken together, these data suggest that at 72 hours post-infection, NTHi still inhibits *Pa* bacterial burden in the lungs. In addition to impeding abundace of *Pa*, sequential infection reduces the severety of damage to airway tissues, as measured by histological scoring, and reduces inflammatory cytokine levels in BALF.

**Figure 6:**
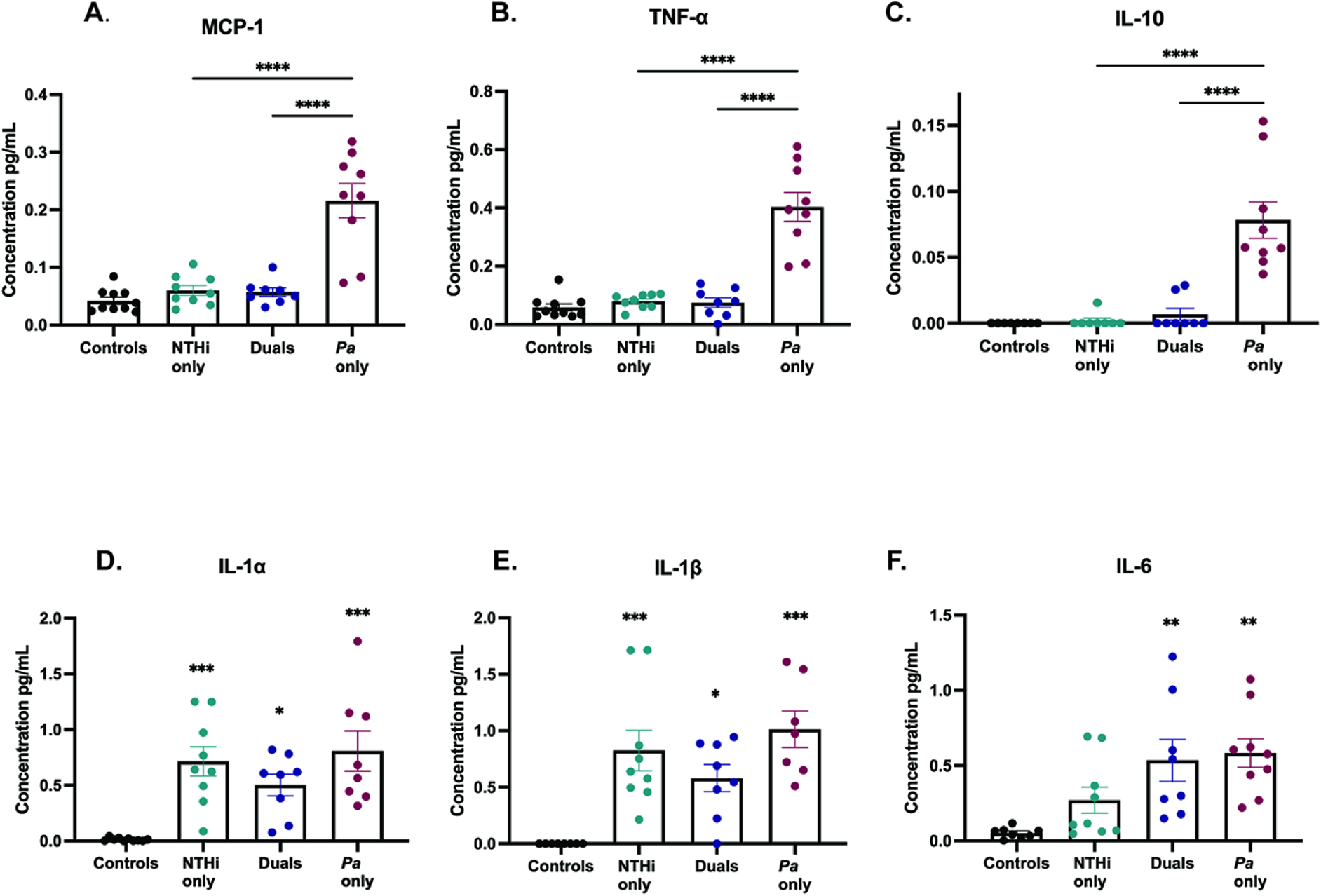
**Inflammatory cytokine levels after dual-infection are reduced compared to *P. aeruginosa* infection alone.** Cytokine and chemokine analyses were performed on BALF supernatant from infected or mock-infected mice 72 hours post-infection using a DuoSet ELISA kits for A) MCP-1, B) TNF-α, C) IL-10, D) IL-1α, E) IL-1β, and F) IL-6. Mean ± SEM, (N = 7-10). Dual infection indicates sequential introduction. One-way ANOVA with Tukey’s multiple comparisons test for post-hoc analysis, *P<0.05, **P<0.01, ***P<0.001, **** P<0.0001.

### Inhibitory effect of nontypeable *H. influenzae* on *P. aeruginosa* infection is independent of bacterial viability

Previous polymicrobial studies involving NTHi suggests that this pathogen may either directly engage in interspecies interaction by active processes, or may be passively involved in priming the host for further immune stimuli (33–36). To determine if live NTHi is required to diminish *Pa* establishment in the lungs, we sequentially infected mice with heat-killed NTHi HI-1r (HK NTHi HI-1r) (inoculum equvialent to ~10^8^ CFU/mL) followed by mucoid *P. aeruginosa* mPA 08-31 (inoculum ~10^7^ CFU/mL) 24 hours later (Fig. S2C). The total infection time of this experiment was 72 hours, repeating our previously established timepoint in which live NTHi infection reduced *Pa* infection. Viable colony counts from lung homogenate showed *Pa* counts were still significantly reduced by pre-treatment with heat-killed NTHi (*P<0.05) (Fig. 7A). Additionally, *Pa* infection resulted in a significantly higher histolopathological score than HK NTHi/*Pa* dual-infection, suggesting that non-viable NTHi is sufficient to protect the lung parenchyma from the severe effects of *Pa* infection, indicating the protection from tissue damage previously seen at this time point during dual-infection is independent of NTHi viability (**P<0.01) (Fig. 7B). H&E stained lung sections harvested from this experiment showed more neutrophilic infiltration and more severe tissue damage after *Pa* single-infection compared to dual-infection (Fig. 7C). As previously seen, the total cell counts in BALF, the percentage of PMN infiltration, and weight loss per animal were not different among single or dual-infection groups, but were significantly increased compared to uninfected controls (*P<0.05, ***P<0.001, ****P<0.0001) (Fig. 5D-E, Fig. S3E). Overall, we found that heat-killed NTHi is sufficient to significantly impede the abundance of *Pa* in the lungs at 72 hours post-infection, suggesting that the interaction between NTHi and *Pa* may be entirely mediated by the immune response of the host.

**Figure 7:**
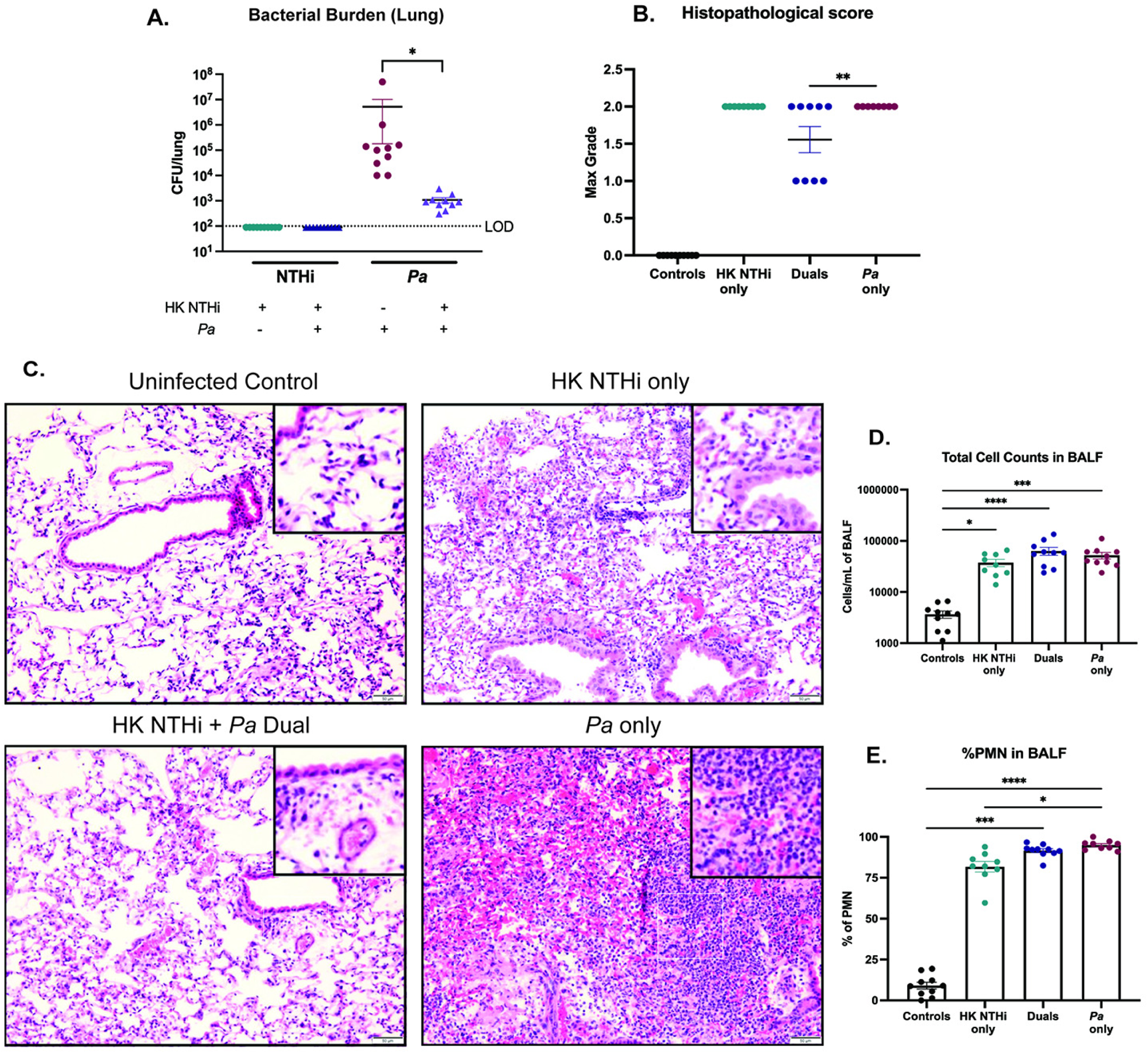
**Impact of nontypeable *H. influenzae* on *P. aeruginosa* infection is independent of bacterial viability.** BALBc/J mice were intratracheally infected with heat-inactivated nontypeable *H. influenzae* (NTHi Hi-1r) 24 hours prior to introduction of mucoid *P. aeruginosa* (mPA 08-31) and assessed 72 hours post-sequential infection. A) Bacterial counts of NTHi and *Pa* in the lung homogenate were enumerated by viable colony counting. (N = 10). Severity of disease was indicated by B) semi-quantitative grading by a pathologist (L.N.) and C) H&E stained lung sections (200x magnification) (N = 8-10). D) Total cell counts in BALF quantified by differential counting. E) Percentage of PMNs in the BALF. (N = 9-10). Mean ± SEM. LOD = limit of detection. Kruskal-Wallis with Dunn’s multiple comparison test, * P<0.05, ** P<0.01, *** P<0.001, **** P<0.0001.

### Sequential introduction with LOS alone reduces the bacterial burden of *P. aeruginosa* in the lungs

Like many mucosal pathogens, NTHi expresses “rough” lipooligosaccharide endotoxins with short non repeating saccharide chains (18). As with other gram-negative endotoxins, NTHi LOS features hexaacylated lipid A which elicits host inflammatory responses via Toll like receptor 4 (19, 20). To test the impact of NTHi endotoxin on *Pa* infection, mice were intratracheally dosed with purified LOS (7 μg/g) 24 hours prior to infection with mPA 08-31, and *Pa* bacterial load from the lung homogenate were assessed 72 hours post infection (Fig. S2C). Bacterial counts revealed that pretreatment with LOS significantly reduces subsequent colonization of *Pa* (***P<0,001) (Fig. 8A). Compared to uninfected controls, purified LOS caused significant weight loss PMN infiltration in BALF (****P<0,0001) (Fig 8B-C), which is consistent with a lung inflammatory response. Based on these findings, we conclude that innate immune priming via NTHi endotoxin is a likely mechanism for the inhibitory effect of NTHi on subsequent *Pa* infection.

**Figure 8:**
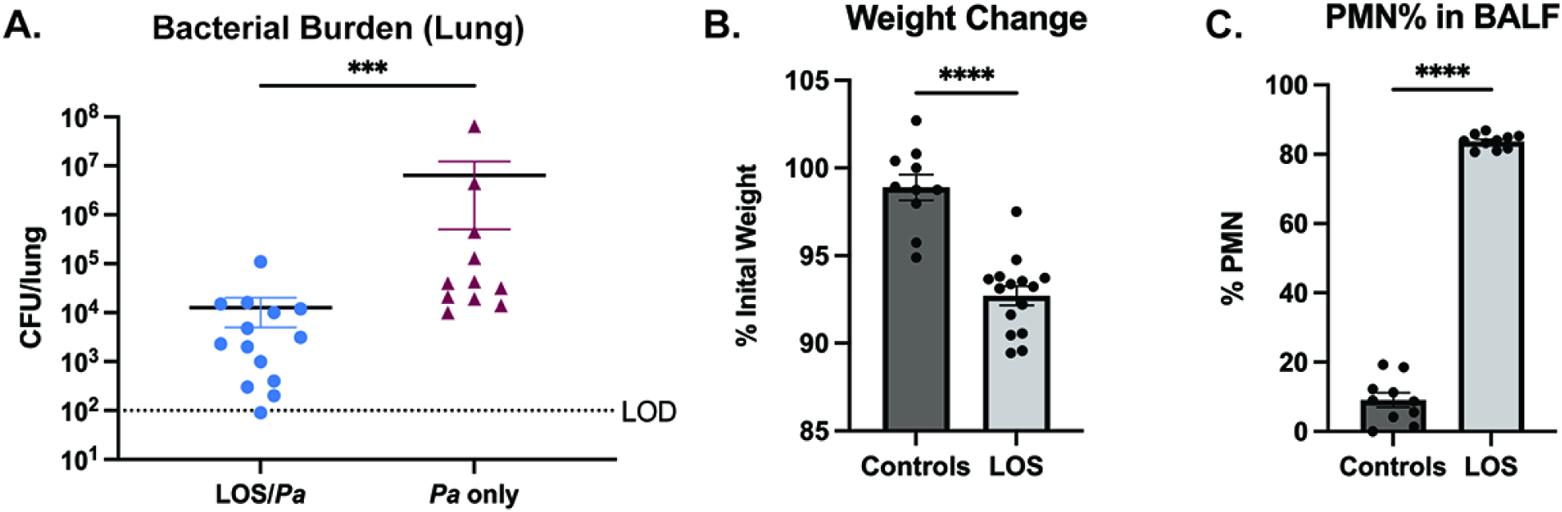
**Challenge with NTHi lipooligosaccharide (LOS) endotoxin elicits host inflammation and reduces colonization of *P. aeruginosa*.** BALBc/J mice were intratracheally dosed with purified LOS (~7 μg/g body weight) 24 hours prior to introduction of mucoid *P. aeruginosa* (mPA 08-31) and assessed 72 hours post-dual challenge. A) Bacterial burden of *Pa* in the lung homogenate with and without prior challenge with LOS. (N = 11-15). LOS alone was sufficient to initiate response to infection, as measured by B) percentage of initial weight and C) percentage of PMNs in BALF, compared to uninfected controls. (N = 10-15). Mean ± SEM. LOD = limit of detection. Mann-Whitney, two-tailed, ***P<0.001, ****P<0.0001.

## DISCUSSION

With the advent and widespread use of culture independent technologies for profiling microbial consortia, it has become clear that the populations within the lungs of patients with CF are diverse and subject to dynamic change. These shifts are likely of importance in determining whether species will persist or be cleared, as well as in the initiation of the inflammatory events that are a hallmark of CF disease (6–8, 27, 37, 38). The consequences of respiratory exacerbations include an intense neutrophilic inflammatory response that can cause significant lung damage and decreased respiratory function, contributing significantly to the morbidity and mortality associated with CF disease (8, 38). There is still much to learn regarding how specific organisms interact within the compromised respiratory tract of CF airways and how these interactions influence disease progression (8).

The complex microbial communities in the lung often change in dominant species and overall population diversity as patients age (22, 27). Decreased microbial diversity and the emergence of mucoid *Pa* as the dominant microbial species are significantly associated with detrimental host immune response, lung functional decline, and eventual patient mortality (22, 27). Previous studies have focused on characterizing key CF pathogens individually, however, these infections are undoubtedly polymicrobial in nature, influencing the pathogenesis of disease as organisms interact (3, 19). Nontypeable *Haemophilus influenzae* is commonly carried within the microbiome of the nasopharynx and upper airways in healthy individuals, and has long been recognized as an opportunistic pathogen in patients with impaired mucociliary defenses including young children with CF (10, 39). In this work, we addressed the role of preceding NTHi infection on susceptibility to respiratory infection with *P. aeruginosa*; the results indicate that NTHi primes an innate host response which impedes *P. aeruginosa* colonization.

Polymicrobial interactions in static biofilms provide basic information about interspecies bacterial interactions. Our results indicate that NTHi and *Pa* form polymicrobial biofilms that support *Pa* growth (Fig. S1, Fig. 1–2). However, to understand polymicrobial infections in patients, we must how these two organisms interact within the lung environment. In a mouse model, we tested how NTHi impacted *Pa* colonization following sequential introduction, representing how patients typically acquire these organisms over time, contrasted with concurrent acquisition.

Bacterial infections in the CF lungs result in chronic inflammation, as microbial and phagocytic debris buildup in the airways, contributing to the reduced lung function and overactive immune response associated with exacerbations (10). The inherently diminished airways defenses associated with CFTR dysfunction are thought to predispose CF patients to frequent microbial infections followed by an inadequate, yet overactive, innate immune response that further initiate tissue damage by airway remodeling (12, 29). Histopathological analysis of lung sections can evaluate immune cell infiltration within alveoli, pleura, perivascular and peri-bronchial spaces, and grade the severity of tissue damage. In CF, the ensuing tissue damage following chronic bacterial infection is a strong indicator of respiratory failure, the leading cause of morbidity and mortality (12). Pre-introduction with NTHi not only reduces the bacterial burden of *Pa*, but also significantly reduces tissue damage compared to a *Pa* single-infection, an effect that may reduce airway remodeling following infection (Fig. 5). While dual-infection reduces the severity of airway parenchymal damage, we wanted to know if inflammatory cytokine levels were reduced after dual-infection as well. The BALF collected from single- and dual-infected mice showed trending decreases in early response cytokines, such as IL-1-α, IL-1-β, and IL-6 in dual-infected mice compared to a *Pa* single-infection (Fig. 6A-C). Since the interleukins are some of the first cytokines to respond to bacterial stimuli, we suspect that significant differences between dual- and single-infected groups may appear at earlier timepoints, prior to 72 hours post-infection (11, 30). We saw a significant decrease in inflammatory cytokine levels of MCP-1, TNF-α, and IL-10 in sequentially infected duals compared to a *Pa* single-infection, suggesting that the reduction in bacterial burden also diminishes a potentially harmful innate immune response (Fig. 6D-F). Most notably, this reduction in TNF-α could extend beneficial effects for CF patients, as the continual production of this early response cytokine indicates an ongoing and overactive innate immune response in the CF lung (11).

Defining how different microbes interact within the CF lung to impact bacterial colonization, persistence, and host response has the potential to guide rational treatment strategies to lengthen periods of asymptomatic or mildly symptomatic infection that are typically associated with younger patients. The results of our infection studies indicate that colonization of the lung with NTHi may provide inherent resistance to subsequent colonization with *Pa*. A likely mechanism for such an effect would be stimulation of innate immune defenses, which can prime host phagocyte responses to respond to incoming pathogens (34, 36, 40, 41). We next addressed the hypothesis that NTHi initiates a host inflammatory response which impedes subsequent *Pa* colonization by testing impact of pretreatment with purified NTHi endotoxin. We thus intratracheally administered purified LOS to mice prior to *Pa* and found that LOS alone promotes the reduction in *Pa* colonization (Fig. 8). These results raise interesting possibilities regarding whether aggressive treatment against early-stage opportunists, such as NTHi, may be contraindicated or even harmful by providing a susceptibility window to other infections. It is also notable that for opportunistic airway infections associated with COPD, an extensive body of recent work indicates that treatment with inflammatory agonists that evoke innate responses can confer protection and are under active investigation as therapeutics (42–46). The concept of therapeutically inducing a low level of airway inflammation to enhance resistance may merit further study in CF related infections.

## MATERIALS AND METHODS

### Bacterial strains and growth conditions

NTHi 86-028NP is well characterized patient isolate for which a fully annotated genome sequence data base is available (47). NTHi HI-1 is a CF patient isolate which was kindly provided by Timothy Starner (University of Iowa Children’s Hospital). NTHi 86-028NPr and NTHi HI-1r were derived from NTHi 86-028NP and NTHi HI-1 by serial passage on plates containing spectinomycin (16). NTHi 86-028NP-gfp+ was constructed by electroporation with plasmid PGM1.1+gfp (provided by K. Mason, Nationwide Childrens’ Research Hospital) (48, 49). For heat-killed NTHi experiments, bacteria were scraped from a plate into sterile PBS, resuspended to ~10^9^ CFU/ml, and incubated at 65°C for 1 h; bacterial killing was confirmed by lack of growth. All NTHi strains were routinely cultured on supplemented brain-heart infusion (sBHI) agar (Difco), containing 10 μg/mL of hemin (ICN Biochemical) and 1 μg/mL of NAD (Sigma). NTHi LOS (a gift from Michael Apicella, University of Iowa) was purified from strain NTHi 2019 according to standard methodology (50) and was intratracheally administered as has been described previously (51, 52).

*P. aeruginosa* mPA08-31 is a mucoid clinical isolate from a CF patient at UAB Hospital, and was provided by S. Birket (University of Alabama at Birmingham) (53, 54). *A* nonmucoid derivative of *P. aeruginosa*, (*P. aeruginosa* FRD1*mucA*+) was provided by J. Scoffield (University of Alabama at Birmingham) (55). *P. aeruginosa* mPA08-31-mCherry+ was derived by electroporation with plasmid pUCP19+mCherry (provided by D. Wozniak, Ohio State University). All *P. aeruginosa* strains were routinely cultured on Luria-Bertani (LB) agar (Difco). Strains were streaked for colony isolation before inoculation into LB broth and shaking overnight at 37°C and 200 rpm.

### Confocal microscopy of static biofilms

*In vitro* biofilms were prepared by resuspending NTHi from an overnight plate culture to~10^8^ CFU/ml in 2x sBHI, or overnight broth culture of *Pa* diluted to ~10^8^ CFU/ml in LB broth. Dual-species biofilms were sequentially seeded with NTHi preceding *Pa* by 12 hours into a 35-mm glass-bottom confocal dish (MatTek) in 2-ml increments such that the final density in each well was ~10^8^ CFU of one or of both organisms. NTHi biofilms were incubated for 12 hours and *Pa* biofilms for 6 hours, respectively. Sequentially seeded polymicrobial biofilms were incubated for 18 hours. Viable colony counts of NTHi or *Pa* from biofilms were obtained by plating on sBHI with spectinomycin or LB, respectively. For fluorescent imaging of NTHi 86028NP-gfp+ and mPA08-31-mCherry+, all dishes were incubated at 37°C with 5% CO_2_ before being washed with sterile PBS and fixed with 4% paraformaldehyde (Alfa Aesar, Tewksbury, MA). Confocal laser scanning microscopy (CLSM) was performed using a Nikon-A1R HD25 confocal laser microscope (Nikon, Tokyo, Japan) at the University of Alabama at Birmingham. All images were processed using the NIS-elements 5.0 software.

### Fluorescent biofilm quantification

Fluorescent biofilms were quantified using BiofilmQ software (56). Single fluorescence channels were automatically segmented using the Otsu algorithm. Background was removed with semi-manually thresholding denoising. Global biofilm parameters of the fluorescence channel representing *Pa* were quantified for both single-species and dual-species biofilms to assess biovolume. 3D spatial representation of cubing (representing 1 μm per cube) was calculated by BiofilmQ and visualized with ParaView according to manufacturer’s instructions.

### Mouse model of respiratory infections

BALB/cJ mice (8 to 10 weeks old) were obtained from Jackson Laboratories (Bar Harbor, ME). For single-species infections, mice were anesthetized with isoflurane and intratracheally infections, mice were anesthetized with isoflurane and infected intratracheally either concurrently with NTHi and *Pa* (~10^9^ CFU NTHi and ~10^8^ CFU *Pa* in 100 μL PBS), or sequentially, in which infection with NTHi occurred 24 hours prior to *Pa* introduction. For viable plate counting, the left lung of each mouse was harvested and homogenized in 500 μL of sterile PBS at 30 Hz/s. Lung homogenate was serially diluted in PBS and plated LB agar to obtain viable colony counts of *Pa* and sBHI agar containing spectinomycin (2 mg/mL) for selection of NTHi HI-1r. All samples from polymicrobial infections were also plated on sBHI without antibiotic for total bacterial counts of both organisms. For histological analysis, the right lung of each animal was inflated with 10% buffered formalin and stored at 4°C until processing. All mouse infection protocols were approved by the University of Alabama at Birmingham (UAB) Institutional Animal Care and Use Committees.

### Bronchoalveolar lavage fluid collection

Bronchoalveolar lavage was performed after euthanasia by flushing mouse lungs with 6 ml of cold PBS in 1-mL increments as previously described (18). Collected BALF was stored on ice until processing and the first 1mL isolated was centrifuged (1350 × *g*, 5 min) to separate the supernatant from immune cells. The cell pellet was used for total and differential cell counts. The remaining 5 mL of collected BALF was stored at −20°C until use for cytokine analyses.

### Cytokine analyses and differential cell counting

Cytokine analyses were performed on the 5 mL of collected BALF via DuoSet ELISA kits (R&D Systems, Minnesota, MN). For total and differential cell counts, cell pellets from BALF were resuspended in fresh PBS collected by cytospin at 500 rpm for 5 minutes. Cells were stained with a Kwik-Diff differential cell stain (ThermoFisher Scientific, Waltham, MA). Three representative fields from each spot were counted to determine the composition of immune cells.

### Histological analysis

Inflated right lungs from infected animals were stored in 10% neutral buffered formalin (Fisher Scientific, Waltham, MA) at 4°C until processing. Sections from each lobe of the right lung were trimmed and sent to the UAB Comparative Pathology Laboratory to be processed, paraffin embedded, sectioned and stained with hematoxylin and eosin (H&E). Images of lung sections (200 x magnification) were taken with Olympus DP25 camera (Tokyo, Japan) using an Olympus BX40 microscope. Semiquantitative grading of all lung sections was performed by a board-certified surgical pathologist (L.N.). Semiquantitative histopathological scores were assigned using a scoring matrix based primarily on neutrophilic influx. Severity was rated on a scale of 0 to 3, where 0 represents no observable neutrophils,1 represents dispersed acute inflammation (mild damage), 2 represents dense infiltration (moderate damage), and 3 represents solid infiltrate with necrosis (severe damage). The final histopathological score represents the maximum grade assigned to each replicate.

### Statistical analyses

All bar graphs represent sample means ± standard error of the mean (SEM). All mouse experiments were repeated at least two times and data from independent experiments were combined for analyses. For nonparametric analyses, differences between groups were analyzed by Kruskal-Wallis test with the uncorrected Dunn’s test for multiple comparisons. For normally distributed data sets, as determined by Shapiro-Wilk normality test, a one-way analysis of variance (ANOVA) was used with Tukey’s multiple-comparison test. Outliers were detected via the ROUT method (Q, 1%) and excluded from the analysis. All statistical tests were performed using GraphPad Prism 9 (San Diego, CA).

## ACKNOWLEDGMENTS

The authors gratefully acknowledge helpful input and discussions with colleagues in the UAB Center for Cystic Fibrosis Research. This work was supported by grants to W.E.S. from the Cystic Fibrosis Foundation (CFFSWORDS1810 and CFFSWORDS20G0). N.L. was a predoctoral trainee supported by the UAB Cystic Fibrosis Foundation Basic Research Center (Steven Rowe, PI). B.C.H. was supported by a fellowship from the UAB Predoctoral Training Program in Lung Biology (T32 HL13640). We thank Michael Apicella (University of Iowa) for providing purified LOS, and Timothy Starner (University of Iowa Children’s Hospital), Daniel Wozniak (Ohio State University), Jessica Scoffield (University of Alabama at Birmingham), and Susan Birket (University of Alabama at Birmingham) for providing bacterial strains.

